# An intracellular pathway controlled by the N-terminus of the pump subunit inhibits the bacterial KdpFABC ion pump in high K^+^ conditions

**DOI:** 10.1101/2021.01.13.426525

**Authors:** Himanshu Khandelia, David Stokes, Bjørn Panyella Pedersen, Vikas Dubey

## Abstract

The heterotetrameric bacterial KdpFABC transmembrane protein complex is an ion channel-pump hybrid that consumes ATP to import K^+^ against its transmembrane chemical potential gradient in low external K^+^ environments. The KdpB ion-pump subunit of KdpFABC is a P-type ATPase, and catalyses ATP hydrolysis. Under high external K^+^ conditions, K^+^ can diffuse into the cells through passive ion channels. KdpFABC must therefore be inhibited in high K^+^ conditions to conserve cellular ATP. Inhibition is thought to occur via unusual phosphorylation of residue Ser162 of the TGES motif of the cytoplasmic A domain. It is proposed that phosphorylation most likely traps KdpB in an inactive E1-P like conformation, but the molecular mechanism of phosphorylation-mediated inhibition and the allosteric links between phosphorylation on the A domain and the inactivation of the pump remain unknown. Here, we employ molecular dynamics (MD) simulations of the dephosphorylated and phosphorylated versions of KdpFABC to demonstrate that phosphorylated KdpB is trapped in a conformation where the ion-binding site is hydrated by an intracellular pathway between transmembrane helices M1 and M2 which opens in response to the rearrangement of cytoplasmic domains resulting from phosphorylation. Cytoplasmic access of water to the ion-binding site is accompanied by a remarkable loss of secondary structure of the KdpB N-terminus and disruption of a key salt bridge between Glu87 in the A domain and Arg212 in the P domain. Our results provide the molecular basis of a unique mechanism of regulation amongst P-type ATPases, and suggest that the N-terminus has a significant role to play in the conformational cycle and regulation of KdpFABC

## 1. Introduction

KdpFABC is a unique K^+^ ion transporting complex expressed in bacteria under low external *K*^+^ concentrations. Bacterial survival under such conditions hinges upon sufficient inward transport of K^+^, and the KdpFABC complex consumes ATP to transport K^+^ against its concentration gradient into the bacterial cytoplasm. The protein thus maintains the requisite high intracellular concentration of K^+^ in low K^+^ growth conditions [1]. Under high external K^+^ conditions, the pump is not required because K^+^ can diffuse into the cell through constitutively expressed K^+^ channels. KdpFABC is a heterotetramer, and the two main functional transport subunits are the ion-channel like subunit KdpA and the pump like subunit KdpB. The KdpA subunit most likely descends from the bacterial KscA family of ion channels, while KdpB clearly belongs to the P-type ATPase family of ion pumps which also include the Na, K ATPase, Ca^2+^-ATPase (SERCA) and the gastric H^+^, K^+^ ATPase [2]. KdpFABC is therefore a one-of-a-kind, unique channel-pump hybrid and an evolutionary outlier. In ion pumps such as SERCA, the ion-binding sites are located in the transmembrane region. The protein undergoes transitions between the so-called E1 and E2 states [3, 4], which face the two different sides of the membrane and have differential affinities for different ions, making the ion-binding site more selective towards the ion present at a lower concentration. The conformational transitions between the E1 and E2 states are driven by the binding of ions to the TM ion-binding sites, and by the ATP-driven phosphorylation and dephosphorylation of the cytoplasmic P domain which contains a canonical aspartate residue which gets phosphorylated. ATP binds to the nucleotide-binding N domain which is an internal kinase and phosphorylates the aspartate residue on the P domain. Finally, the actuator A domain uses a TGES motif to catalyse the dephosphorylation of the aspartate residue in the second half of the cycle [5]. The dephosphorylation and phosphorylation of the P domain are accompanied by significant rearrangements in the relative orientations of the cytoplasmic domains, and this rearrangement drives reorientation of the TM helices. The rearrangement of the TM helices opens intracellular or extracellular pathways leading to the ion-binding sites, through which ions bind, or are released. In this way, ions are transported across the membrane through the TM region. For a more detailed review of the conformational changes which occur during the complete transport cycle of P-type ATPases, the reader can consult several outstanding reviews on the subject [6, 7, 5]. However, the exact mechanism of transport of K^+^ through KdpFABC, and how the KdpA subunit is coupled to the KdpB subunit remains unclear. Crystal [8] and cryo-EM [9] structures show that K^+^ ions bind to the KdpA subunit in its outside open state. It is not clear however, how the external gate of the KdpA subunit closes, or whether the K^+^ ions are transported through the channel subunit, or are transported from the KdpA to the KdpB subunit, and then released into the cytoplasm through the KdpB subunit [10]. The latter scenario is unlikely because in the available high dimensional structures of the pump, there is no conduit of large enough diameter between the KdpA and the KdpB subunit through which a K^+^ ion can travel. Solving the problem of mechanism of TM ion transport probably requires the availability of the complex in conformations different from those observed so far. The focus of this paper is the mechanism of regulation of KdpFABC in high external K^+^ conditions.

In both crystal [8] (Figure 1) and cryo-EM structures [9] of the KdpFABC complex, the Serine residue (Ser162) in the TGES motif is found to be phosphorylated [11], while it is well established that the preceding Glutamate residue (Glu161) catalyses the hydrolysis of the phosphorylated aspartate residue. The role of serine phosphorylation has now been traced to the inhibition of the pump when the cells are transferred to a high K^+^ medium. Under high K^+^ conditions, the KdpFABC pump must be inhibited [12, 13], to avoid ATP hydrolysis and waste of cellular resources when inward ion transport can proceed through passive ion channels without expending energy. Therefore, phosphorylation of the Serine residue (Ser162) is likely to trap the complex into a conformation that can no longer cycle between the E1 and E2 states, thus preventing ATP turnover. The molecular mechanism by which phosphorylation of Ser162 inhibits KdpFABC remains unknown. The Ser162Ala mutation makes the protein constitutively active, while the phosphorylation mimicking mutation Ser162Asp inactivates the protein [11]. Like in the inactivating Glu161Gln mutation, the pump is stuck in an E1-P or E2-P conformation in the Ser162Asp mutation [11]. Most importantly, the pump and channel subunits become uncoupled in either of these mutations, suggesting that phosphorylation (or lack thereof) of Ser162 has long-range allosteric effects that influence the conformational flexibility and transitions of the pump during the transport cycle. Here, we employ Molecular Dynamics (MD) simulations of the unphosphorylated (SER) and phosphorylated (PHOS) Ser162 versions of KdpFABC to show that phosphorylation drives the opening of an intracellular water-filled pathway leading to the KdpB ion-binding sites, through TM helices M1, M2 and M4. The access to this pathway is controlled by the interactions of the N-terminus of KdpB with the A domain. We propose that the opening of this previously unknown intracellular pathway can trap KdpFABC in one of the afore-mentioned phoshorylated EP states, thus applying brakes on the pump catalytic cycle and on ATP hydrolysis.

**Figure 1:**
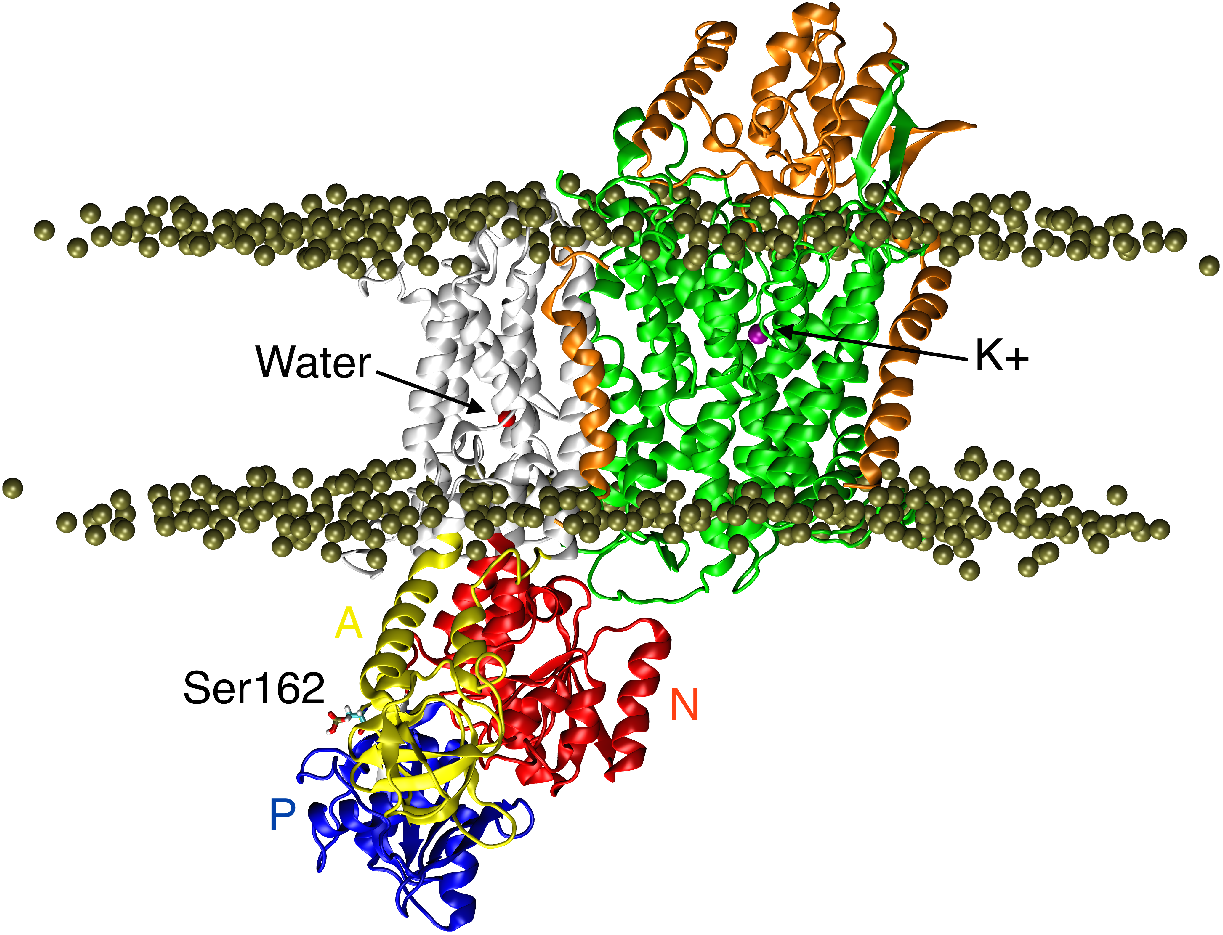
Starting setup of the simulations performed in this work, showing the crystal structure of the KdpFABC heterotetramer complex, pdb id 5MRW. The channel KdpA channel subunit is coloured green while the KdpC and KdpF subunits are orange. A water molecule (red sphere) is visible in the KdpB pump canonical ion-binding site, and a K^+^ ion (purple) is resolved in the KdpA subunit. Residue Ser162 of the TGES motif in the A domain is phosphoryated. Lipid headgroups are coloured tan. For clarity, lipid tails, hydrating water molecules on both sides of the membrane and bulk ions are not shown.

## 2. Results and Discussion

### 2.1. Cytoplasmic domains are highly mobile in phosphorylated Kdp

The simulations reveal distinct and reproducible differences in the behavior of KdpB that are attributable to serine phosphorylation. The first is the mobility of the cytoplasmic domains of KdpB. In particular, the average root mean squared deviations of each cytoplasmic domain relative to their starting positions are significantly higher in PHOS simulations, which can be seen in the RMSD distributions for the individual domains (Figure 2). The *RMSD* fluctuates between 0.2 and 0.6 nm for both systems, suggesting an overall stable tertiary structure in the simulations (Figure 2A).The higher deviations from the starting structure reflect that the protein samples more conformations in the PHOS simulations than in the SER simulations. The possible functional relevance of this is discussed in the following sections. A covariance and principal components analysis further elucidates the differing dynamic behavior in PHOS and SER simulations. The first two eigenvectors of the covariance matrix of atomic fluctuations represent the largest deviations in the dataset (Figure 3) and are used to analyze the global collective motions. For this analysis, each protein conformation sampled during the simulations was projected upon the two dimensional space formed by the first two eigenvectors to obtain a 2D-scatter plot (Figure 3A). In addition, a histogram representing the 1D motion along each eigenvector is shown along the margin of each axis. This projection map reveals several clouds which correspond to distinct protein conformations sampled during the simulations. These clouds are likely to represent free energy minima in the protein conformational landscape which are clearly different for PHOS (blue) and SER (orange) simulations. The analysis is similar to what we performed earlier in the context of a different membrane protein [14]. More specifically, this analysis reveals novel conformations, labeled C and D, that are sampled in the PHOS simulations, whereas the SER simulations remain within the larger cloud of conformations labeled B. A *k-means* clustering algorithm was used to extract representative conformations from each conformational cluster, which are illustrated in panels B, C and D of Figure 3). In conformations C and D, sampled only in the PHOS simulations, the A-domain (yellow) uncouples from the P-domain (blue), and collapses towards the membrane.

**Figure 2:**
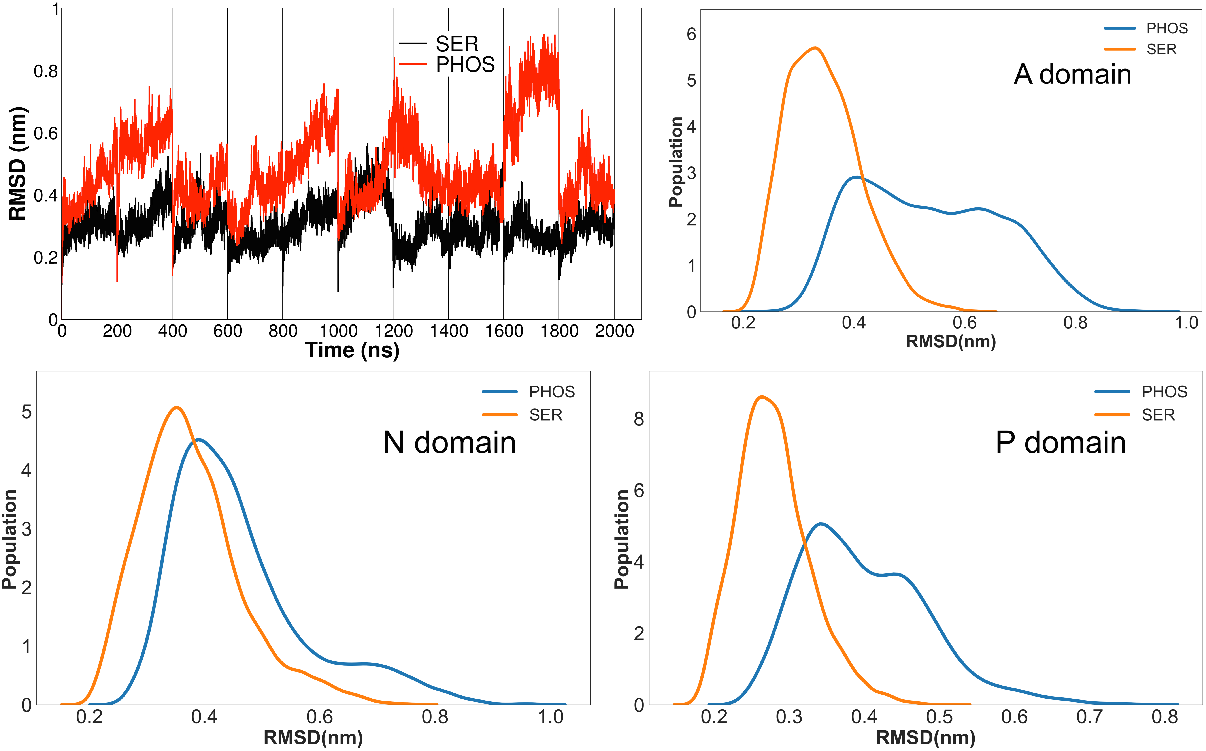
(A) The root mean squared deviation of the cytoplasmic domains of the protein backbone of the PHOS and the SER simulations over 10 different trajectories of 200 ns each. (B) (C) and (D) A histogram of the RMSD of the A, N and P domains. All three cytoplasmic domains deviate more from the starting structure in the PHOS simulations.

**Figure 3:**
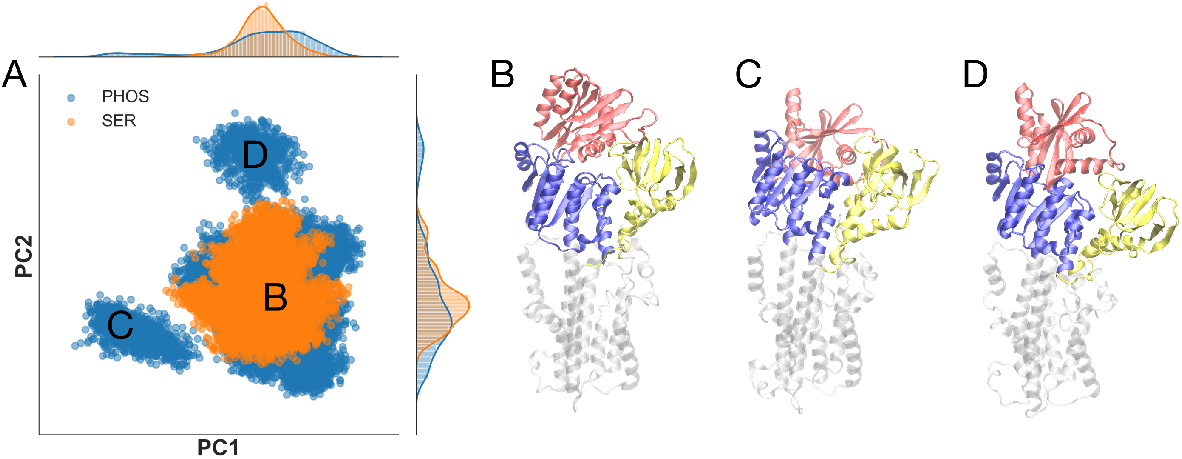
Principle component analysis of conformations sampled during the simulations. (A) Projection of individual frames from the simulation onto the first two eigenvectors (PC1 and PC2) from the covariance matrix. The prevalent conformation represented by the cloud labeled B is sampled by both PHOS and SER simulations. However, the PHOS simulation has sampled novel conformations represented by the peripheral clouds labeled C and D. Projections onto each individual axis are shown along the top and right margins. (B-D) Structures extracted by *k-means* analysis of panel A from clouds B, C and D, respectively. Cytoplasmic domains are colored as follows: A-domain yellow, N-domain red, P-domain blue.

Trajectories projected upon the first eigen vectors for the PHOS and SER simulations are shown in Movies S1 and S2.

### 2.2. Phosphorylation opens an intracellular water pathway leading to the KdpB ion-binding site

The second key difference involves the canonical ion-binding site of P-type ATPases formed near Pro264 on the M4 helix. This site is occupied by a water molecule in the crystal structure of KdpB (Figure 1). In the PHOS simulations, this water molecule escaped almost immediately, whereas it remains bound for the duration of the SER simulations.

This escape is facilitated by the opening of an aqueous channel in KdpB from the cytoplasmic side of the membrane (Figure 4, and supplementary movie S3). As a result, the Pro264-bound water molecule is frequently exchanged with a different water molecule in the aqueous channel. In contrast, water never reaches the canonical binding site and the initial water molecule remained occluded in any of the 10 SER simulations. (Figure 4). To quantify the penetration of water into this site, the radial distribution function g(r), which is a normalised histogram of pair-wise residue distances, is calculated for all water molecules relative to the center of mass defined by Pro264, Thr265, Asp583, and Lys586, which surround the site and coordinate the water molecule in the crystal structure (Figure 4A). These data indicate that this intramembrane site becomes highly hydrated when Ser162 on KdpB is phosphorylated, but not for the unphosphorylated Kdp complex.

**Figure 4:**
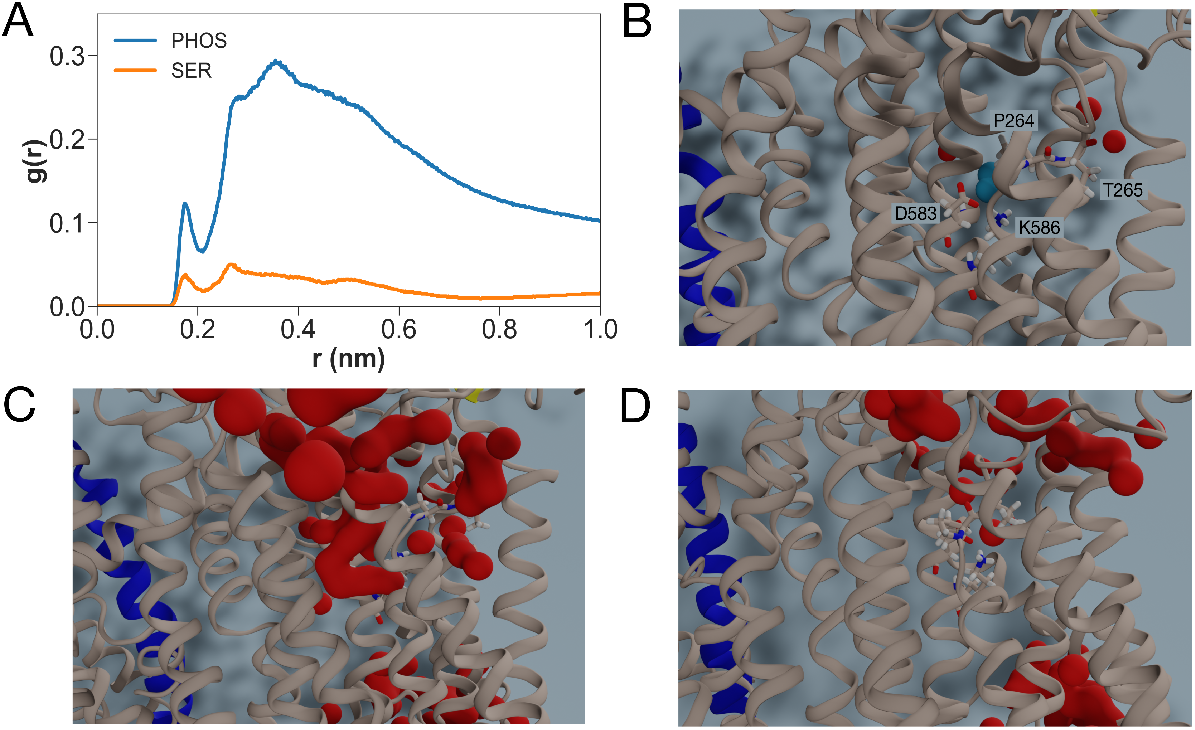
Water accessibility of the canonical ion binding site in KdpB. (A) Radial distribution functions between water and the center of mass of residues Pro264, Thr265, Asp583 and Lys586 from KdpB, which surround the canonical ion binding site. The radial distribution function corresponds to a normalized histogram of distances between two atom selections, averaged over the time of the simulation. (B) Initial configuration of the canonical site for both the SER and PHOS simulations with a water molecule (blue) bound at the location seen in the X-ray structure. (C) and (D) Final frames from PHOS and SER simulations respectively. The canonical ion-binding site is significantly more hydrated in the PHOS simulations with water penetrating the site from the cytoplasmic side of the membrane (top of the panel).

The water pathway opens between helices M1 and M2, and in the case of the SER simulations, the pathway is closed by a salt bridge between Glu87 on the A-domain and Arg212 on the P-domain (Figure 5D). The salt bridge is broken in the PHOS simulations as a result of the large scale motions of the cytoplasmic domains, and in this way separating helices M1 and M2, and opening the water channel. An intracellular pathway between helices M1, M2 and M4 is also postulated for the sarcoplasmic reticulum Ca^2+^-ATPase [15], and was observed in the first crystal structure of SERCA [16], as well as in inhibitor-bound structures of SERCA [17]. The M1 helix probably plays a role in the occlusion of the bound Ca^2+^ in SERCA [18, 19]. Palytoxin, which converts the Na+, K^+^ ATPase to a channel-like protein, opens a continuous ion-pathway which also passes between M1, M2, M4 and M6 on the cytoplasmic side [20]. Whether or not the observed pathway leading to the ion-binding sites in our simulations of the KdpFABC comprise of a functional pathway for K^+^ release into the cytoplasm remains to be seen. However, it is clear that similar pathways are linked to functional roles in other P-type ATPases. In the case of KdpFABC, the pathway could only serve to trap the protein into an inactive state, and not be responsible for ion release into the cytoplasm. The pathway observed in the simulations is not seen in any of the available structures of KdpFABC to date. It is possible that crystal and cryo-EM structures only samples a subset of the large conformational landscape sampled by the phosphorylated pump. So far, no structure of the active dephosphorylated pump has emerged.

**Figure 5:**
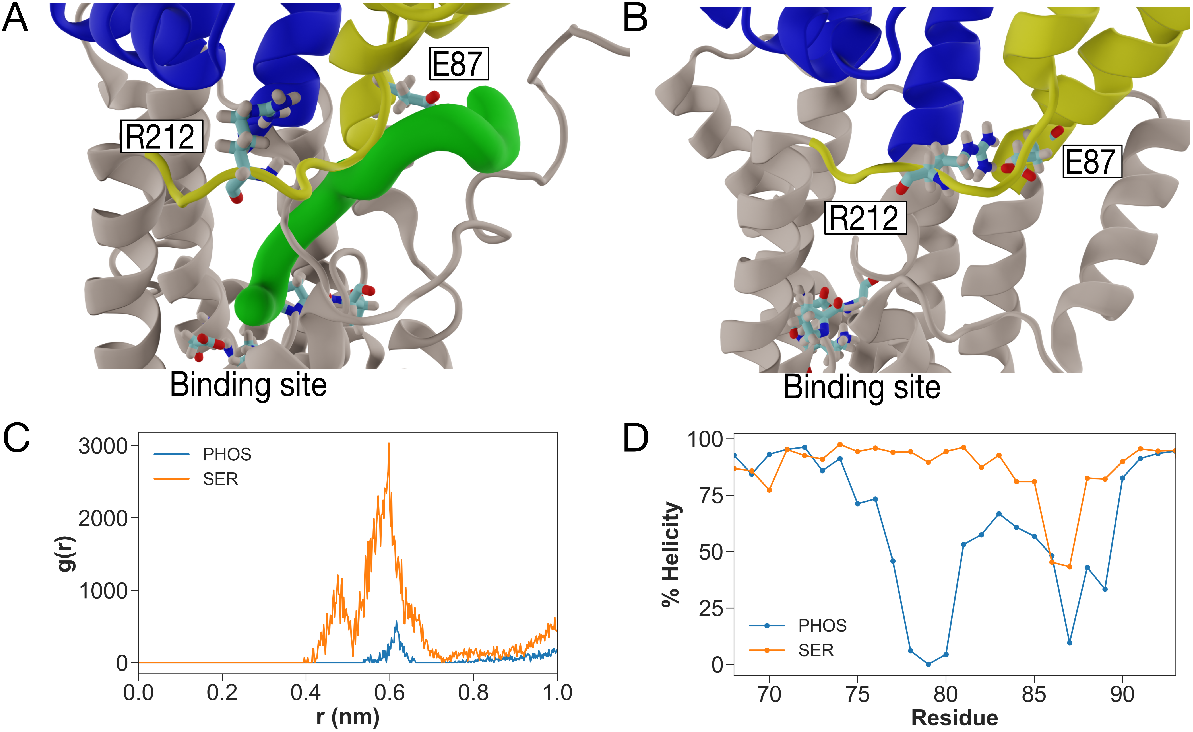
Interactions between M1 and M2 control entry of water into the canonical ion binding site. (A) Snapshot from the PHOS simulations shows that Glu87 and Arg212 are displaced and a water pathway is detected by CAVER (green surface) that leads to the canonical binding site. (B) Snapshot from the SER simulations shows the proximity of Glu87 on M1 and Arg212 on M2. The canonical binding site is below and no water pathway is present. (C) Radial distribution function between the side-chain center of masses of Glu87 and Arg212. Higher peaks for SER simulations indicate stronger interactions between these residues in this system, whereas the lower peaks for PHOS simulations indicate disruption of these interactions. (D) The helical content of the linker between M2 and the A-domain (residues 68-92) is significantly lower for the PHOS simulations indicating unwinding of this structural element.

### 2.3. Hydration of the ion-binding site is accompanied by changes in the conformations of the N-terminus

The opening of the intracellular pathway leading to the ion-binding sites is accompanied by remarkable changes in the secondary structure of the N-terminus of the KdpB. In the SER simulations, residues 11 through 23 remain helical most of the time during the simulations. In the PHOS simulations, however, the helical content of these residues is nearly completely lost (Figure 5D). The conformations of the N-terminus relative to the ion-binding site are shown in (Figure 1). In the SER simulations, the helical N-terminus is linked to the M1 via a random-coil linker comprising of residues 26 through 34, with 27 and 34 being helix-breaking Proline residues. Part of the N-terminus remains membrane-bound, but the rest projects into the cytoplasm to interact closely with residues 83 through 98 on the M2 helix which also protrude into the cytoplasm (Figure 6C). In the PHOS simulations however, the N-terminus is fully unfolded, and collapses parallel to the membrane. In addition, the M2 helix also unfolds near residues 87 through 92, breaking the salt bridge between Glu87 on M2 and Arg212 on top of helix M4. These data are depicted in Figure 6, where it is clear that the distance between the N-terminus and the residues 87 through 92 fluctuates significantly, and is higher in the PHOS simulations, compared to the SER simulations. These data, combined with the earlier claim that the intracellular pathway between helices M1, M2 and M4 could play a functional role, suggest that the N-terminus of KdpFABC has an important role to play in the conformational cycle of the pump, and may even regulate the release of the K^+^ ion into the cytoplasm. The N-terminus was previously shown to regulate the E1-E2 conformational equilibrium of the Na+, K^+^, ATPase by interacting with the cytoplasmic domains [21, 22], and it has since been hypothesized that the N-terminus may play a role in the regulation or the conformations of both the Na+, K^+^, ATPase and the H^+^, K^+^, ATPase [23].

**Figure 6:**
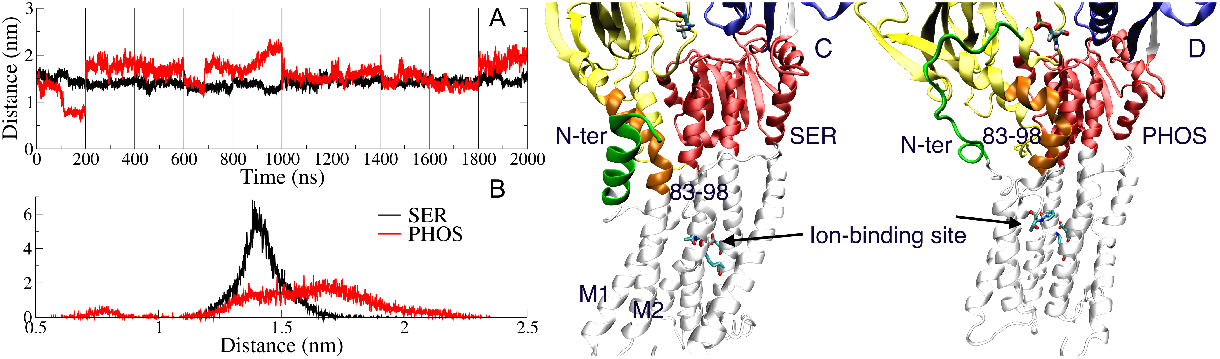
(A) The average distances between the center of mass of the N-terminus (residues 9 through 25) and the cytoplasmic portion of the M2 helix (residus 87 through 92) over all 10 simulations. Except in the first simulation replica, the distance increases and fluctuates more for all PHOS replicas compared to the SER replicas. (B) The histogram of distances shown in A. C and D: The interactions between the N-terminus (green) and residues 83 through 98 (orange) in the A domain in the SER and PHOS simulations respectively. The N-terminus is fully unfolded in the PHOS simulations, but remains helical in the SER simulations. The color coding for the cytoplasmic domains is the same as Figure 1.

Taken together, there are several dynamic phenomena that occur in the PHOS simulations, and which are absent in the SER systems, which lead to the formation of the water pathway leading to the ion-binding sites. These changes are: the unfolding of the N-terminus, the unfolding of the M2 helix between residues 86 and 91, the large-scale movement of the cytoplasmic domains and the broken salt bridge between Glu87 and Arg212. We are reluctant to ascribe any of these as a single potential decisive step which lead to hydration of the binding site. Current models of allostery rather emphasize the dynamic nature of many small conformational changes which add up to altered dynamics of a conformational ensemble of a protein upon binding of an allosteric ligand. We advocate this ensemble view of allostery according to which reactions such as phosphorylation can regulate protein function allosterically without gross conformational changes.

To conclude, we have carried out MD simulations of phosphorylated and dephosphorylated versions of the bacterial KdpFABC complex to propose that the phosphorylation inhibits the protein by opening an intracellular water pathway leading to the ion-binding sites in the KdpB pump subunit. We also hypothesize that the N-terminal residues guard the entrance to the ion-binding site in the absence of Ser162 phosphorylation, and in this way, the KdpFABC N-terminus may have a significant role to play in the regulation of the pump. Since it can control the entry of water (and ions) to the pump subunit ion-binding site, the N-terminus is also likely to be involved in the regulation and kinetics of the overall conformational cycle of the pump. Our hypotheses can be experimentally validated or refuted by simple mutational experiments such as the measurement of pump activity and ATP turnover in a mutant where the N-terminus is truncated. Our simulations are not very long by current standards. However, all simulation replicas point to the same observation with respect to the hydration of the ion-binding site, lending confidence to our sampling protocols. Simulations using the cryo-EM structures as starting models will be implemented in the future, although these structures have a lower resolution than the crystal structure used as a starting model in the current simulations.

## 3. Materials and Methods

To set up the system for simulation, a bilayer containing 550 lipid molecules was constructed using CHARMM-GUI with POPC and POPE in a 4:1 ratio, as a simple mimic of the bacterial membrane. The published crystal structure for KdpFABC (PDB: 5MRW), including the bound potassium ion as well as the protein, was introduced in the bilayer with the help of the OPM server and the system was hydrated with 75,000 water molecules. We used the crystal structure because it had the highest resolution. 150mM of salt (KCl) was added to the aqueous phase. The protonation states of the titratable amino acids were initially set by PROPKA [24]. Several residues had a pKa higher than 7, and were therefore protonated. However, a short 50 ns simulation run showed that all acidic amino acids had a pKa values well below 7.0. With this in mind, we kept all acidic amino acids deprotonated during the production simulations. Ten independent simulations of 200 ns each were launched for KdpFABC with and without serine phosphorylation, referred to as PHOS and SER simulations respectively.

All-atom MD simulations were performed under periodic boundary conditions using GROMACS version 2016.3 [25, 26, 27, 28, 29]. 10 replicates with different initial velocities were simulated for both the PHOS and the SER simulations. We used the TIP3P water model with Lennard-Jones interactions on hydrogen atoms. A 1.2 *nm* cutoff was used for non-bonded neighbour list, and was updated every 10 steps. The van der Waals interactions were switched off between 1.0 to 1.2 *nm*. Electrostatic interactions were treated with the particle mesh Ewald (PME) method [30, 31]. All systems were minimised with 5,000 steps using steepest decent algorithm. For each simulation, the converged system was equilibrated for 5 ns with backbone restraints on the protein followed by a production run of 200 ns. The temperature of the system was kept at 310K with the Nosé-Hoover thermostat [32, 33] after an equilibration run which was performed with the Berendsen thermostat [34]. The pressure was kept at 1 bar with the Parrinello-Rahman barostat [35] along with a semi-isotropic pressure coupling scheme. The Linear Constraint Solver (LINCS) [36] algorithm was used to constrain all bonds containing hydrogen. A 2 fs time step was used and trajectories were sampled every 20 ps. The data analysis was carried out using GROMACS and homemade scripts. The analysis was performed on the concatenated trajectories from the 10 individual simulations. Snapshots were rendered using Visual Molecular Dynamics (VMD)[37]. Data analysis was performed on a single concatenated trajectory for each of the three systems, unless otherwise mentioned. Trajectories for the center of mass of different domains were extracted by MDAnalysis tools [38, 39] and distance and angular data were calculated by homemade scripts.

Covariance and principal components analysis (PCA) was performed using GROMACS. After removing center of mass translation and rotation, the covariance analysis was performed for the protein backbone atoms. First, a covariance matrix *C* is extracted from atomic fluctuations

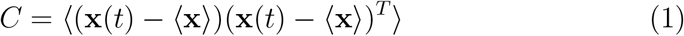

where x is a 3*N* -dimensional column vector describing the coordinates of the *N* protein backbone atoms, and x(*t*) are the positions at time *t*. The triangular brackets indicate an ensemble average. *C* is a 3*N* x 3*N* symmetric matrix, and is then diagonalised by a coordinate tranformation *R*

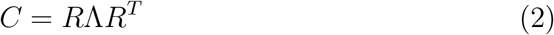

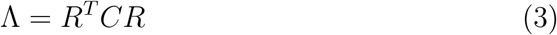

This orthogonal transformation transforms *C* into a diagonal matrix Λ = *diag*(*λ*_1_, *λ*_2_, …*λ*_3*N*_), which contains the eigenvalues *λ*_*i*_ of the covariance matrix *C*. The *i*th column of *R* contains the *i*th eigenvector **r**_*i*_ with the corresponding eigenvalue *λ*_*i*_.

The global collective motions of the trajectory can then be obtained by projecting the trajectory ensemble onto individual eigenvectors, to obtain the principal components **p**_*i*_, *i* = 1, 2, …*N* by taking an internal product between the transpose of the eigenvector and the atomic fluctuation.

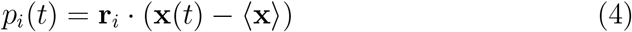

Each trajectory snapshot, thus projected on two different eigenvectors, yields one point on the point cloud distributions shown in Figure 3.

## 4. Acknowledgements

HK and VD are funded by Lundbeckfonden grant numbers R82-2011-7280. The simulations were performed on the ABACUS 2.0 supercomputer at the Danish e-Science Center at SDU, the Piz Daint supercomputer under grant number pr98 and on the Kay supercomputer under the PRACE DECI grant number ANNEX. The financial support by the Novo Nordisk Foundation grant (number NF18OC0032608) for the ROBUST-project is also acknowledged. DS acknowledges funding from the NIH grant number R01 GM108043

The authors thank Wojciech Kopec for useful discussions.

## References

[1] A. Ballal, B. Basu, S. K. Apte, The kdp-atpase system and its regulation, J Biosci 32 (2007) 559–68. URL: https://www.ncbi.nlm.nih.gov/pubmed/17536175. doi:10.1007/s12038-007-0055-7.

[2] M. Bramkamp, K. Altendorf, J.-C. Greie, Common patterns and unique features of p-type ATPases: a comparative view on the KdpFABC complex fromEscherichia coli(review), Molecular Membrane Biology 24 (2007) 375–386. URL: https://doi.org/10.1080/09687680701418931. xdoi:10.1080/09687680701418931.

[3] R. L. Post, S. Kume, T. Tobin, B. Orcutt, A. K. Sen, Flexibility of an active center in sodium-plus-potassium adenosine triphosphatase, Journal of General Physiology 54 (1969) 306–326. URL: https://doi.org/10.1085/jgp.54.1.306. xdoi:10.1085/jgp.54.1.306.

[4] R. W. Albers, Biochemical aspects of active transport, Annual Review of Biochemistry 36 (1967) 727–756. URL: https://doi.org/10.1146/annurev.bi.36.070167.003455. xdoi:10.1146/annurev.bi.36.070167.003455.

[5] J. V. MÃÿller, C. Olesen, A. M. Winther, P. Nissen, The sarcoplasmic ca2+-atpase: design of a perfect chemi-osmotic pump, Quarterly reviews of biophysics 43 (2010) 501–66. doi:10.1017/S003358351000017X.

[6] M. G. Palmgren, P. Nissen, P-type atpases, Annual review of biophysics 40 (2011) 243–66. URL: http://www.ncbi.nlm.nih.gov/pubmed/21351879. xdoi:10.1146/annurev.biophys.093008.131331.

[7] C. Toyoshima, R. Kanai, F. Cornelius, First crystal structures of na+, k+-ATPase: New light on the oldest ion pump, Structure 19 (2011) 1732–1738. URL: https://doi.org/10.1016/j.str.2011.10.016. xdoi:10.1016/j.str.2011.10.016.

[8] C. S. Huang, B. P. Pedersen, D. L. Stokes, Crystal structure of the potassium-importing kdpfabc membrane complex, Nature 546 (2017) 681–685. URL: https://www.ncbi.nlm.nih.gov/pubmed/28636601. xdoi:10.1038/nature22970.

[9] C. Stock, L. Hielkema, I. Tascon, D. Wunnicke, G. T. Oostergetel, M. Azkargorta, C. Paulino, I. Hanelt, Cryo-em structures of kdpfabc suggest a k(+) transport mechanism via two inter-subunit half-channels, Nat Commun 9 (2018) 4971. URL: https://www.ncbi.nlm.nih.gov/pubmed/30478378. xdoi:10.1038/s41467-018-07319-2.

[10] B. P. Pedersen, D. L. Stokes, H.-J. Apell, The KdpFABC complex – k+ transport against all odds, Molecular Membrane Biology 35 (2019) 21–38. URL: https://doi.org/10.1080/09687688.2019.1638977. xdoi:10.1080/09687688.2019.1638977.

[11] M. E. Sweet, X. Zhang, H. Erdjument-Bromage, V. Dubey, H. Khandelia, T. A. Neubert, B. P. Pedersen, D. L. Stokes, Serine phosphorylation regulates the p-type potassium pump KdpFABC, eLife 9 (2020). URL: https://doi.org/10.7554/elife.55480. xdoi:10.7554/elife.55480.

[12] D. B. Rhoads, L. Laimins, W. Epstein, Functional organization of the kdp genes of escherichia coli k-12, J Bacteriol 135 (1978) 445–52. URL: https://www.ncbi.nlm.nih.gov/pubmed/355227.

[13] A. J. Roe, D. McLaggan, C. P. O’Byrne, I. R. Booth, Rapid inactivation of the escherichia coli kdp k+ uptake system by high potassium concentrations, Mol Microbiol 35 (2000) 1235–43. URL: https://www.ncbi.nlm.nih.gov/pubmed/10712703. doi:10.1046/j.1365-2958.2000.01793.x.

[14] V. Dubey, B. Bozorg, D. Wüstner, H. Khandelia, Cholesterol binding to the sterol-sensing region of niemann pick c1 protein confines dynamics of its n-terminal domain, PLOS Computational Biology 16 (2020) e1007554. URL: https://doi.org/10.1371/journal.pcbi.1007554. doi:10.1371/journal.pcbi.1007554.

[15] M. Bublitz, M. Musgaard, H. Poulsen, L. Thøgersen, C. Olesen, B. Schiøtt, J. P. Morth, J. V. Møller, P. Nissen, Ion pathways in the sarcoplasmic reticulum ca2+-ATPase, Journal of Biological Chemistry 288 (2013) 10759–10765. URL: https://doi.org/10.1074/jbc.r112.436550. doi:10.1074/jbc.r112.436550.

[16] C. Toyoshima, M. Nakasako, H. Nomura, H. Ogawa, Crystal structure of the calcium pump of sarcoplasmic reticulum at 2.6 {aa resolution, Nature 405 (2000) 647–655. URL: https://doi.org/10.1038/35015017. doi:10.1038/35015017.

[17] M. Takahashi, Y. Kondou, C. Toyoshima, Interdomain communication in calcium pump as revealed in the crystal structures with transmembrane inhibitors, Proceedings of the National Academy of Sciences 104 (2007) 5800–5805. URL: https://doi.org/10.1073/pnas.0700979104. doi:10.1073/pnas.0700979104.

[18] A. P. Einholm, J. P. Andersen, B. Vilsen, Roles of transmembrane segment m1 of na+, k+-ATPase and ca2+-ATPase, the gatekeeper and the pivot, Journal of Bioenergetics and Biomembranes 39 (2007) 357–366. URL: https://doi.org/10.1007/s10863-007-9106-x. doi:10.1007/s10863-007-9106-x.

[19] M. M. G. Geurts, J. D. Clausen, B. Arnou, C. Montigny, G. Lenoir, R. A. Corey, C. Jaxel, J. V. Møller, P. Nissen, J. P. Andersen, M. le Maire, M. Bublitz, The SERCA residue glu340 mediates interdomain communication that guides ca2+ transport, Proceedings of the National Academy of Sciences (2020) 202014896. URL: https://doi.org/10.1073/pnas.2014896117. doi:10.1073/pnas.2014896117.

[20] A. Takeuchi, N. Reyes, P. Artigas, D. C. Gadsby, The ion pathway through the opened na+, k+-ATPase pump, Nature 456 (2008) 413–416. URL: https://doi.org/10.1038/nature07350. doi:10.1038/nature07350.

[21] N. Boxenbaum, S. E. Daly, Z. Z. Javaid, L. K. Lane, R. Blostein, Changes in steady-state conformational equilibrium resulting from cytoplasmic mutations of the na, k-ATPase α-subunit, Journal of Biological Chemistry 273 (1998) 23086–23092. URL: https://doi.org/10.1074/jbc.273.36.23086. doi:10.1074/jbc.273.36.23086.

[22] Q. Jiang, A. Garcia, M. Han, F. Cornelius, H.-J. Apell, H. Khandelia, R. J. Clarke, Electrostatic stabilization plays a central role in autoinhibitory regulation of the na +, k + -ATPase, Biophysical Journal 112 (2017) 288–299. URL: https://doi.org/10.1016/j.bpj.2016.12.008. doi:10.1016/j.bpj.2016.12.008.

[23] D. Diaz, R. J. Clarke, Evolutionary analysis of the lysine-rich n-terminal cytoplasmic domains of the gastric h+, k+-ATPase and the na+, k+-ATPase, The Journal of Membrane Biology 251 (2018) 653–666. URL: https://doi.org/10.1007/s00232-018-0043-x. doi:10.1007/s00232-018-0043-x.

[24] C. R. Søndergaard, M. H. M. Olsson, M. Rostkowski, J. H. Jensen, Improved treatment of ligands and coupling effects in empirical calculation and rationalization of pKa values, Journal of Chemical Theory and Computation 7 (2011) 2284–2295. URL: https://doi.org/10.1021/ct200133y. doi:10.1021/ct200133y.

[25] H. J. Berendsen, D. van der Spoel, R. van Drunen, Gromacs: a messagepassing parallel molecular dynamics implementation, Computer physics communications 91 (1995) 43–56.

[26] D. Van Der Spoel, E. Lindahl, B. Hess, G. Groenhof, A. E. Mark, H. J. Berendsen, Gromacs: fast, flexible, and free, Journal of computational chemistry 26 (2005) 1701–1718.

[27] B. Hess, C. Kutzner, D. Van Der Spoel, E. Lindahl, Gromacs 4: algorithms for highly efficient, load-balanced, and scalable molecular simulation, Journal of chemical theory and computation 4 (2008) 435–447.

[28] S. Pronk, S. Páll, R. Schulz, P. Larsson, P. Bjelkmar, R. Apostolov, M. R. Shirts, J. C. Smith, P. M. Kasson, D. Van Der Spoel, B. Hess, E. Lindahl, Gromacs 4.5: a high-throughput and highly parallel open source molecular simulation toolkit, Bioinformatics 29 (2013) 845–854.

[29] M. J. Abraham, T. Murtola, R. Schulz, S. Páll, J. C. Smith, B. Hess, E. Lindahl, Gromacs: High performance molecular simulations through multi-level parallelism from laptops to supercomputers, SoftwareX 1 (2015) 19–25.

[30] T. Darden, D. York, L. Pedersen, Particle mesh ewald: An n.log(n) method for ewald sums in large systems, The Journal of chemical physics 98 (1993) 10089–10092.

[31] U. Essmann, L. Perera, M. L. Berkowitz, T. Darden, H. Lee, L. G. Pedersen, A smooth particle mesh ewald method, The Journal of chemical physics 103 (1995) 8577–8593.

[32] S. Nosé, A unified formulation of the constant temperature molecular dynamics methods, The Journal of chemical physics 81 (1984) 511–519.

[33] W. G. Hoover, Canonical dynamics: Equilibrium phase-space distributions, Phys. Rev. A 31 (1985) 1695–1697.

[34] H. J. Berendsen, J. v. Postma, W. F. van Gunsteren, A. DiNola, J. Haak, Molecular dynamics with coupling to an external bath, The Journal of chemical physics 81 (1984) 3684–3690.

[35] M. Parrinello, A. Rahman, Polymorphic transitions in single crystals: A new molecular dynamics method, Journal of Applied physics 52 (1981) 7182–7190.

[36] B. Hess, H. Bekker, H. J. Berendsen, J. G. Fraaije, Lincs: a linear constraint solver for molecular simulations, Journal of computational chemistry 18 (1997) 1463–1472.

[37] W. Humphrey, A. Dalke, K. Schulten, Vmd: visual molecular dynamics, Journal of molecular graphics 14 (1996) 33–38.

[38] R. J. Gowers, M. Linke, J. Barnoud, T. J. Reddy, M. N. Melo, S. L. Seyler, D. L. Dotson, J. Domanski, S. Buchoux, I. M. Kenney, O. Beckstein, Mdanalysis: a python package for the rapid analysis of molecular dynamics simulations, in: Proceedings of the 15th Python in Science Conference, volume 98, 2016.

[39] N. Michaud-Agrawal, E. J. Denning, T. B. Woolf, O. Beckstein, Mdanalysis: A toolkit for the analysis of molecular dynamics simulations, Journal of Computational Chemistry 32 (2011) 2319–2327.

